# What is the cost of that fence? The impact of fences on the movements of ungulates in a hyper-arid landscape

**DOI:** 10.64898/2026.02.10.704417

**Authors:** Morgan L. Hauptfleisch, Stefanie Urban, Lindesay Scott-Hayward, Monique Mackenzie, Kenneth Uiseb

## Abstract

Ungulate movements in arid environments are largely driven by rain events, food resources and surface water availability. In hyper arid areas such as the Namib desert these are patchily distributed, fluctuating and overall sparse. As a result, animals living in these environments need to be highly mobile to exploit the ephemeral and spatiotemporally variable resources. In the past few decades, there has been growing recognition of the importance of wildlife habitat connectivity, and the detrimental effects of linear infrastructure on wildlife and their movements. Barriers, such as roads and fences, block or filter wildlife movements, with severe and sometimes lethal effects on wildlife especially in dry periods or resource-poor environments. In the Greater Sossusvlei Namib Landscape we assessed whether fences impacted ungulate home ranges and movements, and identified particular sections of fences or roads which were most restrictive to ungulate movements. To achieve this, the movements of 12 springbok (*Antidorcas marsupialis*), 13 gemsbok (*Oryx gazella*) and 15 Hartmann’s mountain zebra (*Equus zebra hartmannae*) were tracked telemetrically. In general, ungulate home range sizes were smaller in the vicinity of physical barriers. Roads and fences were found to impact ungulate movements considerably in some areas: these included the C14 and C19 main roads that run from the coast to Maltahöhe and from Solitaire to Maltahöhe respectively, several district roads, parts of the Namib-Naukluft National Park fence, as well as farm fences. While Hartmann’s mountain zebra were able to cross some fences, springbok and gemsbok were not as successful, their movements sometimes being completely restricted within farms or along fences until they found a fence gap to cross. The findings highlight which barriers are key to consider for modification to allow for wildlife movement.

## Introduction

The Greater Sossusvlei-Namib Landscape (GSNL), located in the south-west of Namibia, is one of five Protected Landscape Conservation Areas in Namibia, which were established under the Namibia Protected Landscape Conservation Areas Initiative (NAMPLACE). The GSNL encompasses the state-owned Namib-Naukluft National Park (NNP) at its core and the adjacent freehold farms and private nature reserves. One objective of the GSNL is to create a fence-free Namib and thereby facilitate the reestablishment of age-old wildlife movement between the Namib desert and the Great Western Escarpment, which lies to its east (Odendaal & Shaw, 2010). This is currently pursued by the active removal of fences or by not repairing the fences which are damaged (Tarr, 2014). The goal is to enable wildlife movements in response to rainfall events and food availability. A baseline survey carried out by NAMPLACE in 2013 showed that 52% of farms in the GSNL were informally connected to the NNP due to removed or damaged fences (Tarr, 2014). Despite this, many fences remain as barriers to wildlife movement, since they are needed for livestock and game farming, or are required by law, e.g. for private landowners to hunt or trade with wildlife adequate fencing is required under the Nature Conservation Ordinance 5 of 1975.

When rainfall in the Namib is low or non-existent, wildlife need to move towards to escarpment in the east and when there is good rainfall, they are attracted to the palatable grass of the Namib plains in the west (Odendaal & Shaw, 2010). Freedom to move in response to rainfall patterns will become increasingly important in the light of climate change and the predicted increase in surface temperatures (Engelbrecht et al., 2024). Wildlife will respond by moving their distribution along climatic gradients (Schloss et al., 2012), such as tracking shifts in rainfall and temperatures (Nunez et al., 2019), and beyond currently established protected areas (Hering et al., 2022, 2023). An animal’s movement is linked to fitness outcomes (Liedvogel et al., 2013) in response to short-term goals, such as gaining energy, reproduction, and survival, including escaping predators and avoiding unfavourable environmental conditions, as well as long-term goals, such as inbreeding avoidance and population survival (Holyoak et al., 2008). This is particularly important in arid environments where unpredictable and patchy rainfall results in a constantly changing amount and distribution of resources for the survival of wildlife (Hauptfleisch et al., 2024).

While the economic (especially agricultural) and ecological uses and benefits of fences are clear in some cases, there are several costs, either intended or unintended, associated with fences (Smith et al., 2020). These effects are experienced by both target species (i.e. those species whose movement fences are intended to restrict) and non-target species. Common intended effects of fences include separation (e.g. to prevent disease spread), exclusion of predators and pests, and redirection (e.g. of wildlife towards over- or underpasses), while unintended effects are numerous, see Smith et al. (2020) for a global review, and Lindsey et al. (2012) for a review of Africa. Fences have several direct and indirect ecological effects (Jakes et al., 2018) which are either immediate or occur over long time spans (Gadd, 2012). Direct negative impacts include injury and mortality due to entanglement of wildlife in fence wires (Rey et al., 2012). Indirect effects on wildlife result in alterations to biology and behaviour at the individual or population level. If not planned according to conservation objectives, fences can fragment and isolate wildlife populations, thereby constitute a barrier to gene flow, and resulting in inbreeding, reduced individual and population fitness, and possibly population extinction (Hilty et al., 2012; Jaeger & Fahrig, 2004; Smith et al., 2020; White et al., 2018). Fenced-off areas can also lead to increased population density and overpopulation of species (Slotow, 2012; Welch & Parker, 2016), with negative effects on resource availability as well as land degradation (Fryxell & Sinclair, 1988), resulting in a decrease in carrying capacity and population declines, if not managed correctly (Moseby et al., 2020; Welch & Parker, 2016). Fences can lead to changes in animal distribution, impacting the competitive balance of species in a guild and thus affecting community structures (Cozzi et al., 2013). Fencing off parcels of land in heterogenous landscapes decreases the variety of resources and thus the range of options that are available to herbivores, hence affecting herbivores in low quality or resource poor patches negatively and decreasing the carrying capacity of land (Boone & Hobbs, 2004).

Although there has been some removal of fences within the GSNL, we hypothesize that much of the required range of large ungulates in this hyper-arid environment is still fragmented by fencing and possibly roads. The aim of this study was therefore to use existing telemetric movement data to determine the effect of fences and roads on the movements of collared springbok (*Antidorcas marsupialis*), gemsbok (*Oryx gazella*) and Hartmann’s mountain zebra (*Equus zebra hartmannae*) in the eastern Namib Naukluft Park and adjacent privately owned land. These species are the main large ungulates resident in this hyper-arid environment. We further sought to identify fences and roads which presented the most resistance to movement and prioritise them for possible removal or modification to allow for movement of wildlife.

## Methodology

### Study Area

The Greater Sossusvlei-Namib Landscape is located in the Hardap and Erongo regions of Namibia (Figure *1*) and includes the large 49,768 km^2^ Namib-Naukluft National Park (NNP) at its core as well as a number of neighbouring farming and tourism establishments. The landscape focal area, which encompasses the landholdings adjoining the NNP and is the area where most of the collared animals occur, is wedged between the Great Western Escarpment to the east and the Namib Sand Sea to the west, in the so-called Pro-Namib (Coetzee, 1971; Odendaal & Shaw, 2010). The landholdings constitute various types of land uses, including tourism, commercial livestock farming (both cattle and small-stock), and game farming, as well as private reserves, including the Tsondab Valley Scenic Reserve to the north and the NamibRand Nature Reserve in the south of the GSNL. Since the study area is approximately 260 km (north-south) by 80 km (east-west) results are spatially illustrated in three regions north, mid, and south to allow adequate scale to be depicted in map results.

**Figure 1.**
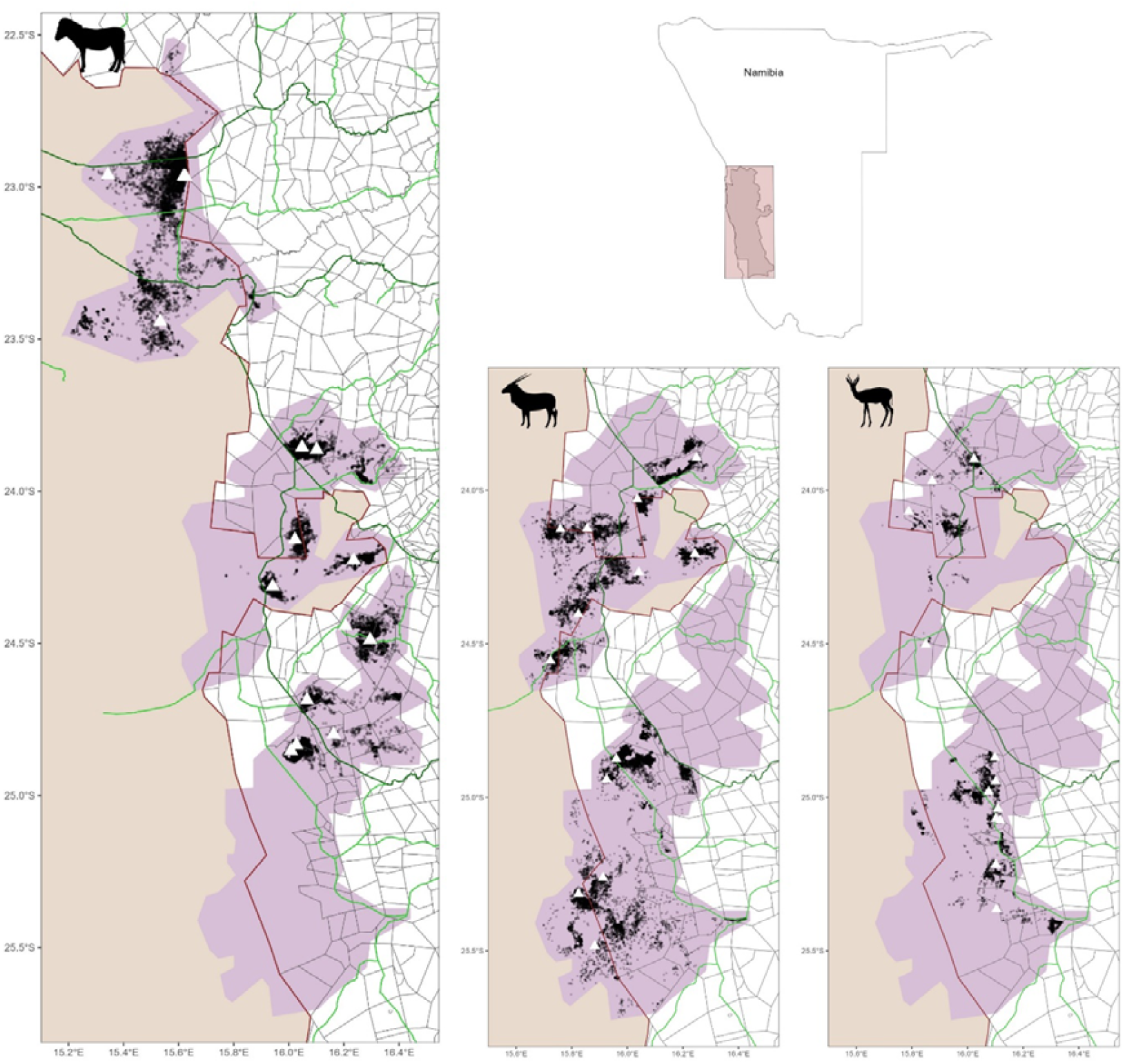
The Greater Sossusvlei Namib Landscape depicting the three regions of the study area (purple), the Namib-Naukluft National Park (beige with red fence line) and the GPS locations of all collard animals. The white triangles are the initial tagging location for each animal. Property fences are shown in grey and major and minor roads in dark and light green.

### Data collection

Satellite collars were fitted to12 springbok (*Antidorcas marsupialis*), 13 gemsbok (*Oryx gazella*) and 15 Hartmann’s zebra (*Equus zebra hartmannae*) in the Greater Sossusvlei-Namib (**Error! Reference source not found**.). The deployment of collars was performed by a wildlife veterinarian from the Ministry of Environment Forestry and Tourism (MEFT) as part of their management mandate to reduce human-wildlife conflict. Only gemsbok females were collared, to attempt the tracking of herds and not just individuals. However, both Hartmann’s zebra males and females were collared as males do not display territoriality. Even though springbok males can be territorial, mostly males were collared as they are physically more robust and are less likely to be negatively affected by the burden of capture and being collared. Only one springbok female was collared. The choice of individuals was based on size and body condition of the animals. Hence, only fully-grown, adult animals were collared. Although not an ideal situation, these factors, combined with difficulty to access many parts of the landscape, and the infrequency of sightings of the species in the large landscape led to the collaring being less selective than if only one sex was collared for comparison. However, since the movement patterns following resources are expected to outweigh social and territorial behaviour in this hyper-arid landscape, the selection of individuals was considered adequate.

**Table 1:** Summary of collared animals in the Greater Sossusvlei-Namib Landscape.

| Species | No. of males | No. of females | Total number | Tracking duration (days) | No. of points |
| --- | --- | --- | --- | --- | --- |
| Springbok | 11 | 1 | 12 | 3124 | 11242 |
| Gemsbok | 0 | 13 | 13 | 9106 | 27526 |
| Hartmann's zebra | 10 | 5 | 15 | 8452 | 31331 |

Based on research by Stiegler et al. (2024) the first two weeks of post capture data were removed from the data sets, in order to take into account the effects of capturing, handling and collaring on animal behaviour and post capture movement rates.

### Homerange analysis

We used a spatially adaptive Binomial Generalised Additive Model (GAM) for home range analysis fitted using the R package MRSea v1.6 (Scott-Hayward et al., 2024). This method was used to account for the residual autocorrelation often present in sequential location data (Estevinho Santos Faustino, 2020) and permits spatially adaptive smoothing, which is particularly valuable for surfaces with uneven and patchy structures (Scott-Hayward et al., 2015, 2024, 2025; Walker et al., 2011).

In these models, flexible univariate relationships were permitted for determining distance to nearest barrier alongside a spatially adaptive two-dimensional flexible surface based on spatial coordinates to accommodate additional patterns. The flexibility of the 1D and 2D smooths was chosen using a Spatially Adaptive Local Smoothing Algorithm (Scott-Hayward et al., 2015, 2025; Walker et al., 2011) with the Bayesian Information Criterion (Schwarz, 1978) as the objective fitness measure. As the location points were correlated in time (the covariates are unlikely to explain this correlation in full), a panel structure (year-month) was used to calculate robust standard errors to accommodate the residual correlation (Scott-Hayward et al., 2015). The location data for each tag (presences) were turned into presence-absence data by randomly selecting absence locations from the bounding box of the tag locations plus 2 km in every direction. These pseudo-absences were selected at a ratio of 5 absences : 1 presence.

Predictions were made to a 500m x 500m grid in the region specific to each tag. Owing to the pseudo-absence generation, the predictions are relative probabilities of presence. Home range was defined as the grid cells that encompass the top 90% of the relative probabilities of presences. The uncertainty around these predictions was estimated using a parametric bootstrap (1000 replicates) and underpinned by the robust variance-covariance matrix. The bootstrap predictions were then used to get 95 percentile-based confidence intervals for the home range size.

### Predictive modelling data preparation

Three distinct regions of data were identified and labelled as North, Mid and South (Figure 1). The data set was split by species and for each one, rasterised using a 500m x 500m grid covering the three regions. In each cell, the number of tag locations and distance to the nearest barrier was recorded.

Barriers are defined as farm fences and major and minor roads.

### Statistical modelling

The modelling methodology is based on the premise that the distribution of each animal, for each species, is driven by a complex mix of environmental factors alongside some possible inhibition by one or more of the fences or roads between farms (barriers). For this reason, we have focused our analysis on quantifying the effects of any fences, using models which explicitly partition any barrier effects from any underlying environmental factors which might be driving their distribution spatially. Specifically, we estimate the barrier effects by fitting a model with terms for both environmental and barrier-related effects and then compare the geo-referenced predictions of this model with geo-referenced predictions based on the scenario where no barrier related-effects are explicitly present. In this alternative reality, the distribution patterns are solely driven by a complex mix of environmental factors alone, which we capture flexibly using a spatially adaptive two-dimensional smoother.

It must be noted, that these modelling results are effectively ‘data-driven’ and this approach does not impose or induce any kind of barrier related effects. Indeed, if no such signal exists in the data then the model will reflect this by evidencing no statistically significant differences in the geo-referenced predictions with and without any barrier related effects. This approach also permits the magnitude of any barrier related effects to be quantified and accompanied by confidence intervals which are adjusted for the longitudinal nature of the data. This provides some context to the extent of any barrier related effects and if they pose a serious block to movement.

A Poisson-based spatially adaptive GAM was fitted for each species, across each of the regions. The first model had two covariates: distance to the nearest barrier and a two-dimensional smooth of coordinates. The barrier relationship was permitted to be flexible via a B-spline while two-dimensional smooth term was fitted using the Complex Region Spatial Smoother (CReSS) to permit a spatially adaptive surface (Scott-Hayward et al., 2015, 2025).

More specifically, the response, (*y*) was the number of tag locations per cell with a Quasi-Poisson distribution which assumes a more flexible mean-variance relationship compared with a traditional Poisson model. The flexibility of the smooth terms was determined using the Spatially Adaptive Local Smoothing Algorithm (Scott-Hayward et al., 2015, 2025; Walker et al., 2011) with the QBIC to determine ‘best fit’. All models were fitted using the R package MRSea v1.6 (Scott-Hayward et al., 2024) and diagnostics for each model were carried out to assess the validity of each. Geo-referenced predictions were then created based on a 500m x 500m grid and uncertainty estimated using a parametric bootstrap of the model coefficients (500 replicates) and the robust variance-covariance matrix.

For each species, we also fitted a second, simpler model which only included a re-fitted spatial term and omitted any fence-related terms. Crucially, in this model we did not re-estimate the flexibility of the spatial term (and only updated the coefficients) to prevent the spatial term adapting to capture the fence related e□ects. Based on this simpler model, we created a second set of geo-referenced predictions with associated confidence intervals, and to enable us to identify important barrier locations we calculated the difference between the predictions under the full and simpler model and built 95% confidence intervals for these differences. Maps were then created with the geo referenced estimated number of tag locations (from the full model) overlaid with crosses indicating grid cells with significantly more locations than expected (as determined by confidence intervals not including zero).

## Results

### Home ranges

Collared animals ranged in NamibRand Nature Reserve, as well as in the Namib-Naukluft National Park to the west and the north, and privately owned farmland of the Rant, Naukluft and Tsaris mountains to the east of the GSNL. A comparison of estimated home ranges of collared individuals shows that there is great variability in home range sizes between species, as well as between individuals of each species (Figure 2). Gemsbok and particularly Hartmann’s zebra home range sizes were more variable compared to springbok home range sizes (Figure 2). In general, springbok (n = 12) had the smallest mean estimated home rangesof 140 km^2^ (±96) with a mean core area of 43 km^2^ (±29). This was followed by Hartmann’s zebra (n = 15) with a mean estimated home range size of 236 km^2^ (±240) and mean core area of 74 km^2^ (±79). Gemsbok (n=13) had the largest mean estimated home range size of 282 km^2^ (±289), which was almost double the size of springbok home ranges, and a mean core area of 86 km^2^ (±81). Gemsbok SAT1770 in the southern region and Hartmann’s zebra SAT1092 in the north region had markedly large home ranges (1,120 km^2^ and 925 km^2^ respectively).

**Figure 2.**
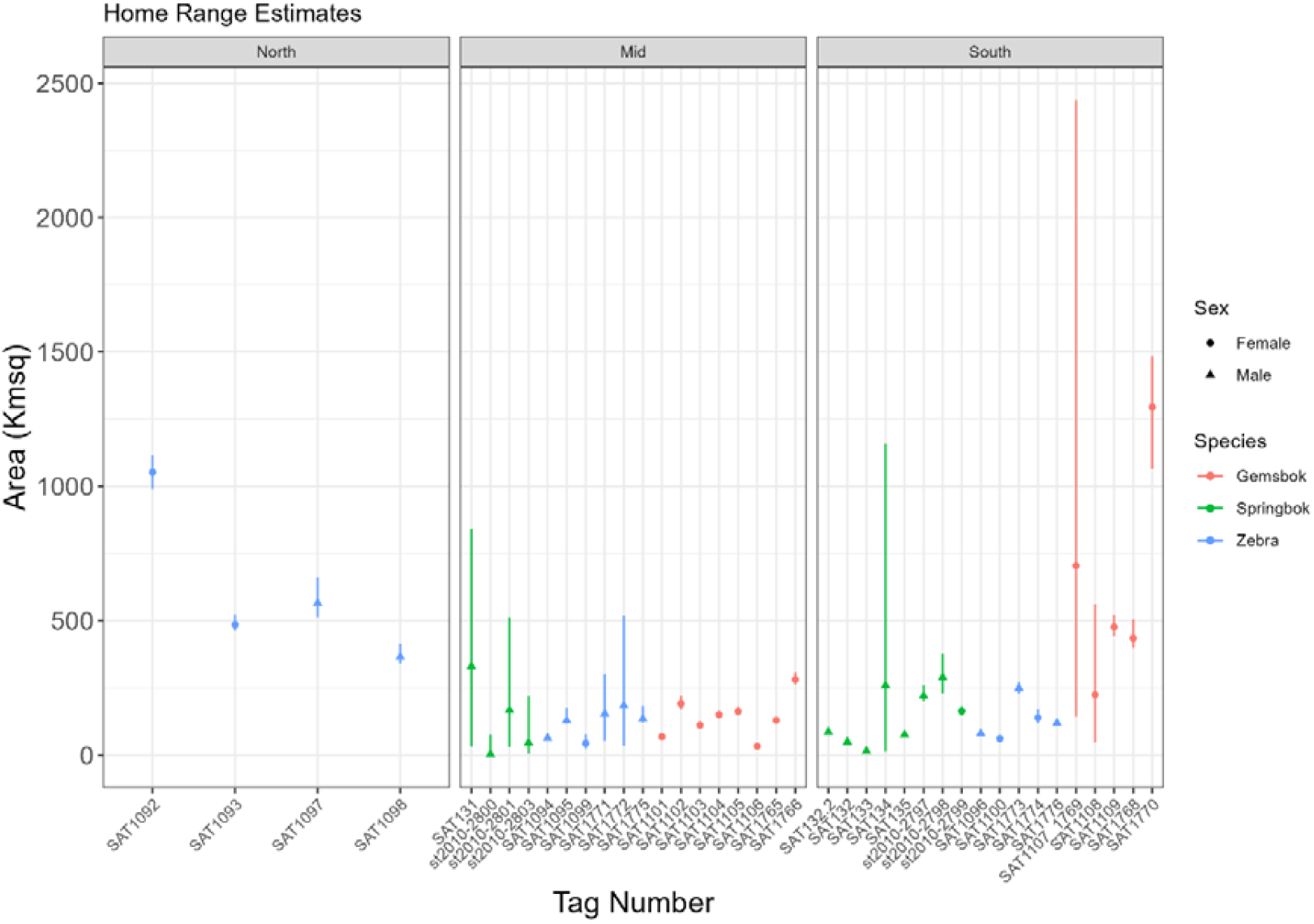
Homerange sizes per individual per species across the Greater Sossusvlei Namib Landscape.

### Wildlife proximity to barriers

The full models estimated the universal relationship between the expected number of telemetry locations and distance to a barrier (road/fence). There was a general trend of more Hartmann’s zebra telemetry locations being close to any barrier (road or fence), with fewer points far from barriers (Figure 3a). Gemsbok showed a general trend of being located at a distance of approximately 3 to 5 km from any barrier (Figure 3b). There is a general trend that springbok locations were often close (less than 1 km) to barriers or 3 to 5 km from barriers (Figure 3c).

Realistically, these estimated relationships are not consistent for all barriers and in some areas, there are more animals than expected near a barrier. This could be because the barrier is causing a blockage and animals are up against the fence, or that the fence is porous and not a barrier at all.

**Figure 3.**
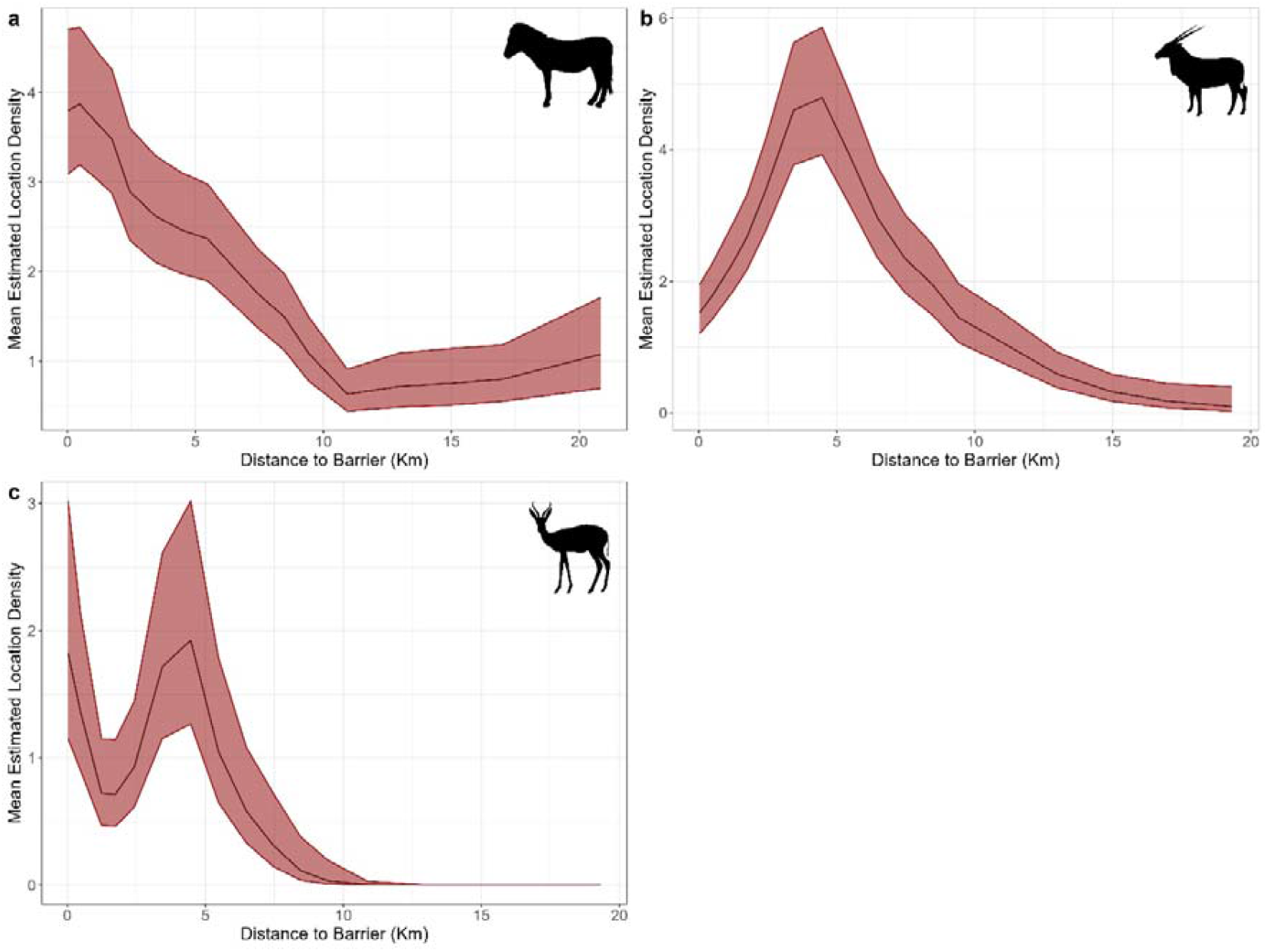
Relationship between distance to nearest barrier and the density of Hartmann’s zebra locations (a), gemsbok (b) and Springbok (c), with upper and lower 95% confidence interval.

### Identifying specific fence pressure areas

The next sections identify specific barriers for each species and region where telemetry locations were higher than expected under the universal barrier effect. The outputs show the area the tracking data covers in pale blue. The ‘+’ symbol on the plots (Figures 4 to 10) indicates where there are significantly more animals within 1km of the barrier than would be expected under the universal barrier relationship shown above. To give context to the porosity of the barriers, the crossing points from the tracking data are shown in yellow.

#### Hartmann’s mountain zebra

There was a significantly high proportion of Hartmann’s zebra locations close to the NNP fence (Figure 4) between the C28 and D1962. The two roads are important crossing points for free movement evidenced by the large number of crossings made. However, the movements of animals onto farmland to the east were clearly impacted by the NNP fence. More locations than expected were found near the fence and no crossings were made.

In the mid region, the C14 main road and where it meets the C19 were identified as a significant barrier to movement of Hartmann’s zebra. Several farms to the north-east of the C14 and the eastern corner of the NNP boundary were shown to have porous boundaries but with significantly more tag locations than expected near these boundaries (Figure 5).

In the southern region, the D0850 and neighbouring farm fences to the north and south had a disproportionately high distribution of location points, however, there was relatively free movement from north to south across the D0850 (Figure 6). The east and west ends of this road were found to be less porous and potential hard barriers to movement. In the south-west of this region, the C19 was shown to be a significant barrier to movement with a larger than expected number of location points close by and no observed crossings.

**Figure 4.**
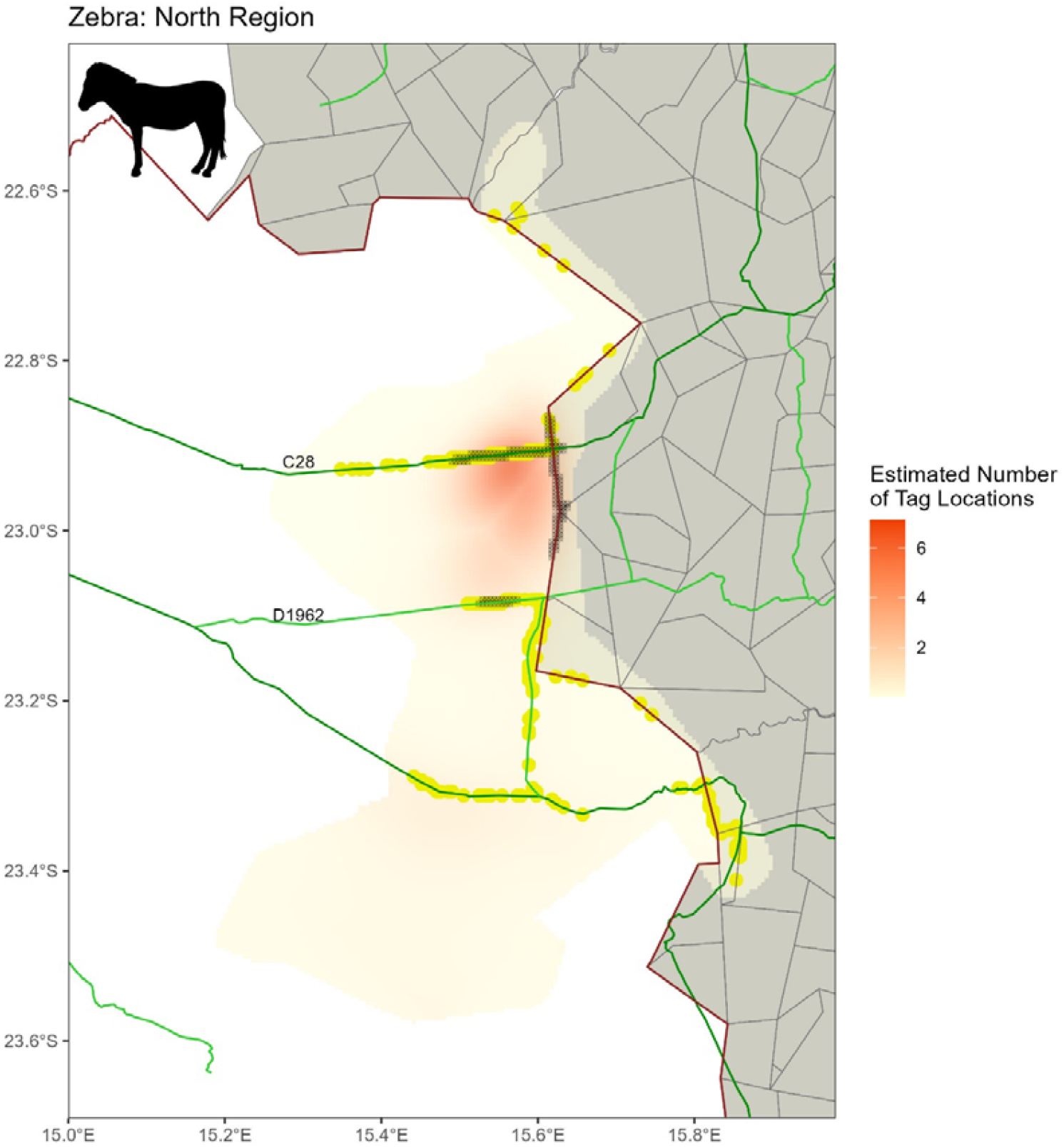
Estimated number of Hartmann’s zebra telemetry locations per cell. The ‘+’ symbols on the plots indicate where there are significantly more telemetry locations within 1 km of the barrier than would be expected under the universal barrier relationship. Grey lines: farm boundary, red line: NNP boundary, dark green lines: main roads, lime green lines: district roads.

**Figure 5.**
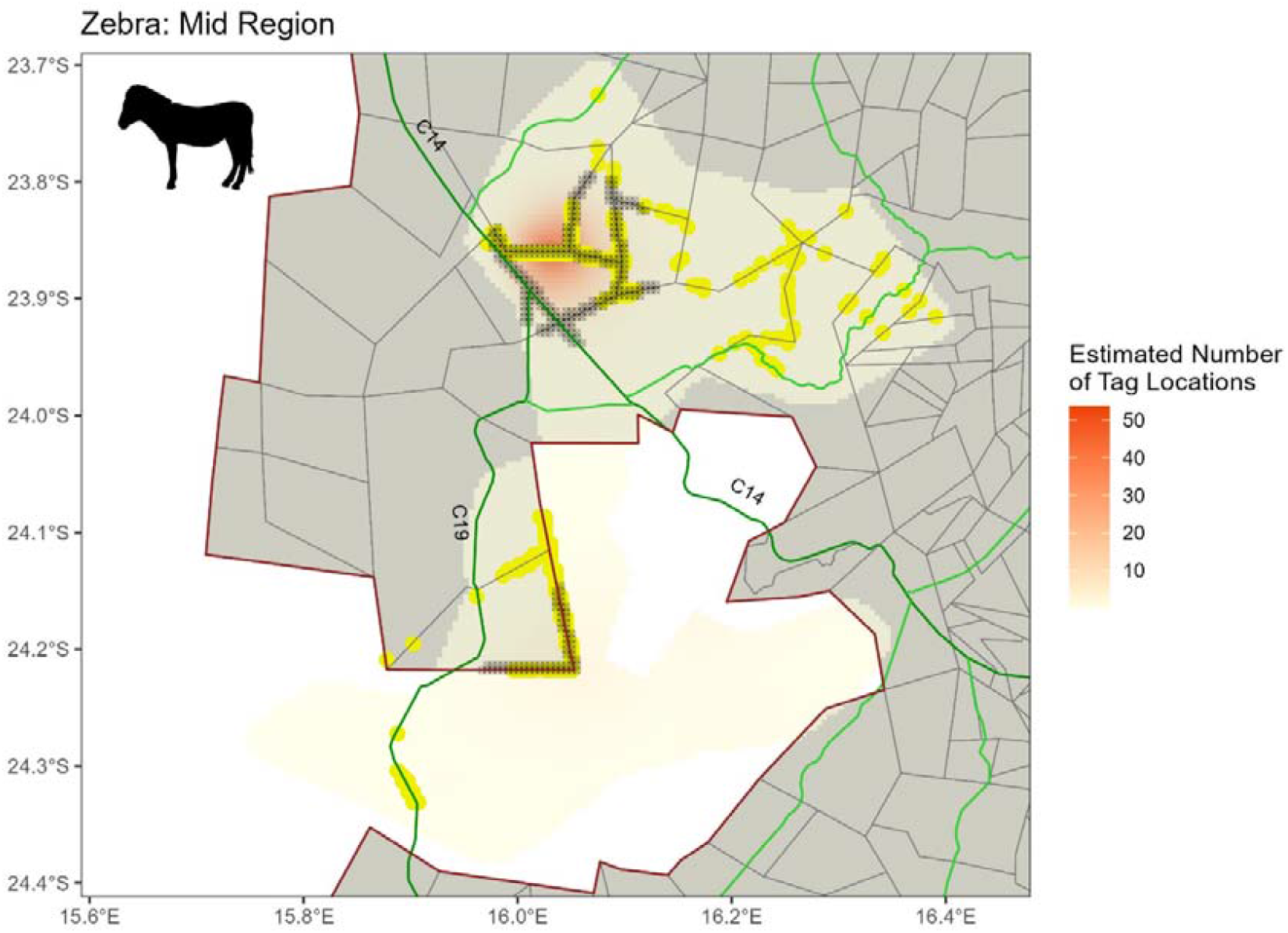
Estimated number of Hartmann’s zebra telemetry locations per cell. The ‘+’ symbols on the plots indicate where there are significantly more telemetry locations within 1 km of the barrier than would be expected under the universal barrier relationship. Grey lines: farm boundary, red line: NNP boundary, dark green lines: main roads, lime green lines: district roads.

**Figure 6.**
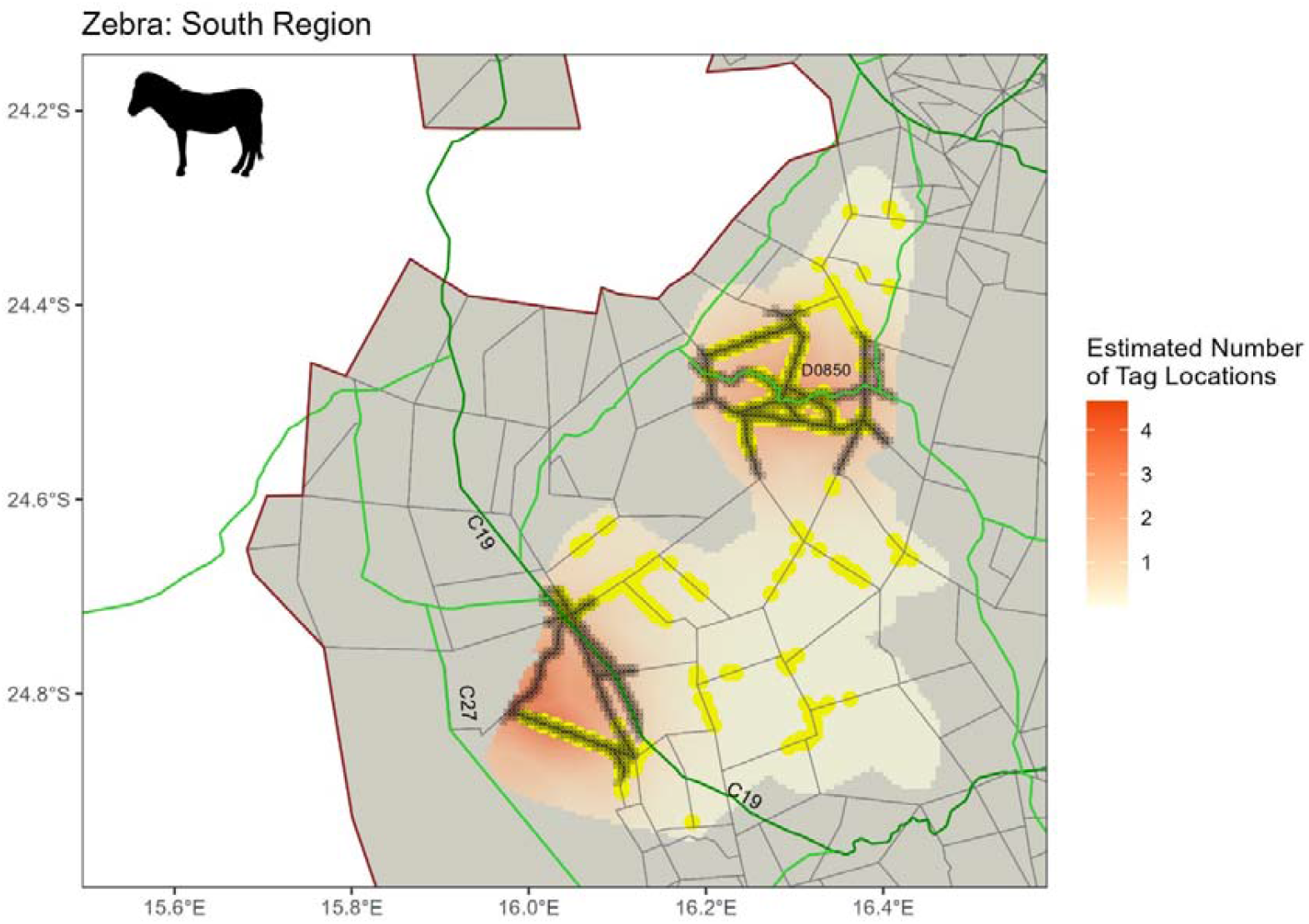
Estimated number of Hartmann’s zebra telemetry locations per cell. The ‘+’ symbols on the plots indicate where there are significantly more telemetry locations within 1 km of the barrier than would be expected under the universal barrier relationship.

#### Gemsbok

In the mid region, the highest number of tag locations of gemsbok were in the north-east. In this area, the C24 district road and where it intersects the C14 main road were found to be causing a barrier to free-movement (Figure 7). The C19 cuts across the NNP and more locations than expected are estimated close by. However, with the exception of the southern section, there are a large number of crossings and no barrier to east-west movement.

Most of the NNP fence that was covered by tag locations was considered important for movement with a large number of crossings observed (Figure 7). A few sections of this fence, particularly the north-east section of the tongue and the triangular section just north of the C27 in the south are more solid and restrict movements. Within the part of the NNP which intrudes as a tongue into the farmland (grey area), there is a north-east to south-west fence (livestock farm Weltevrede) and section of the C19 within this farm which have a higher-than-expected estimated number of locations and no crossing suggesting a well-maintained fence forming a solid barrier to movement. The C27, NNP and farm fences in the southern portion of the mid region have a higher-than-expected number of locations near these barriers than would be expected without them, but do not cause significant inhibition of movement with large numbers of crossings evidenced.

In the southern region, the area close to the C19 road showed a significant increase in the number of gemsbok locations than would be present without this barrier (Figure 8). The figure shows minimal crossing points and no tags were able to move substantially eastward across the road. There were also a number of farm fences south of this road causing significant movement barriers. The NNP fence with the NamibRand Nature Reserve is porous in places however, it has a larger than expected number of locations close to the fence indicating its importance in east-west movement.

**Figure 7.**
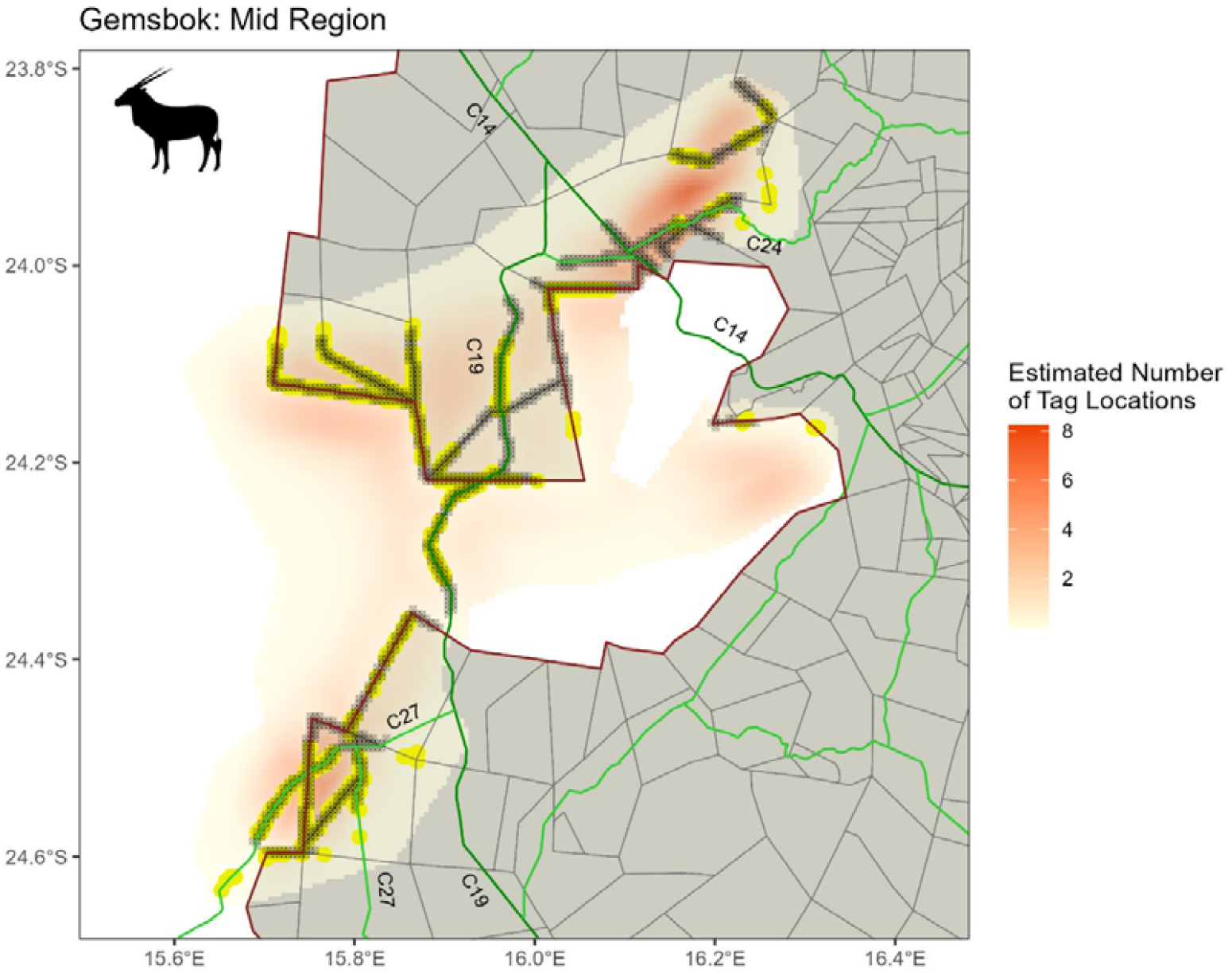
Estimated number of Gemsbok telemetry locations per cell. The ‘+’ symbols on the plots indicate where there are significantly more telemetry locations within 1 km of the barrier than would be expected under the universal barrier relationship Grey lines: farm boundary, red line: NNP boundary, dark green lines: main roads, lime green lines: district roads.

**Figure 8.**
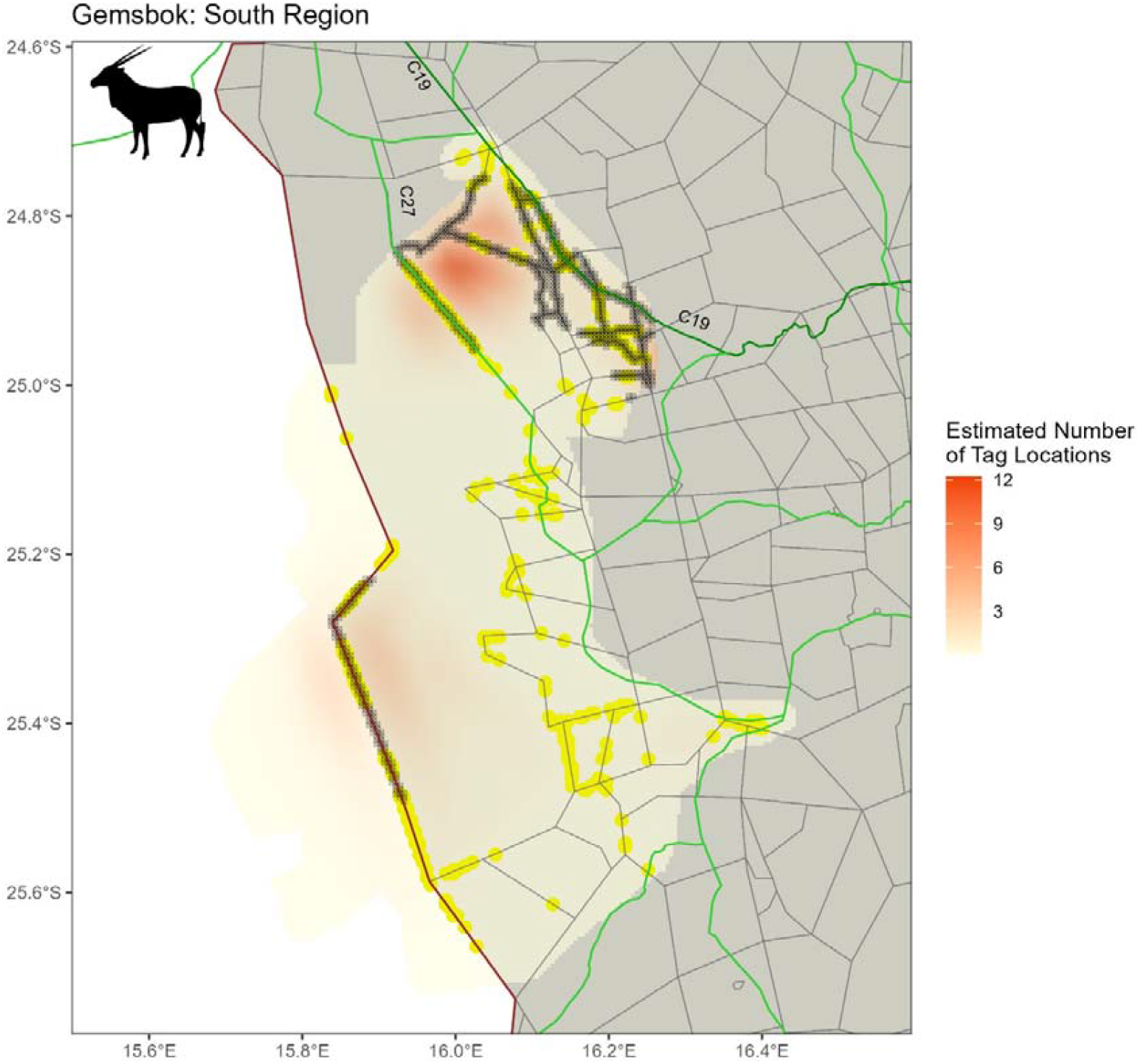
Estimated number of Gemsbok telemetry locations per cell. The ‘+’ symbols on the plots indicate where there are significantly more telemetry locations within 1 km of the barrier than would be expected under the universal barrier relationship. Grey lines: farm boundary, red line: NNP boundary, dark green lines: main roads, lime green lines: district roads.

#### Springbok

There was a significantly higher number of springbok location points within 1 km of the C14 and C19 main roads (Figure 9). Some blockage was caused by the C14 road (although springbok were able to cross at a number of points along the road). The C19 was marked as a potential obstruction from the C14 down to the NNP fence. At the joining of the C14 and the middle of the NNP tongue, springbok were able to cross, however, there is large section in the middle which poses an obstruction to springbok movement.

Like for gemsbok, the Weltevrede livestock farm boundary (north-east to south-west fence in the tongue), restricted north-south springbok movement despite some crossings possible) (Figure 9). However, unlike Hartman’s zebra or gemsbok, springbok were not able to easily cross the NNP fence line. There were more locations than expected estimated in the south-west and west sections of the tongue and no crossings suggesting a barrier to movement.

In the southern region of the study area, there were more springbok locations than expected where the C27 crosses the NamibRand and east of this where the NamibRand borders farmland (Figure 10).

The boundary fence was a solid barrier to movement with no crossings. Some crossings of the C27 were evident however this was patchy indicating a potential obstruction. Further south, where the C27 travels through farmland (grey area in Figure 10) there were also significantly more springbok locations than expected and no crossing points making this road a significant barrier to movement (Figure 10).

**Figure 9.**
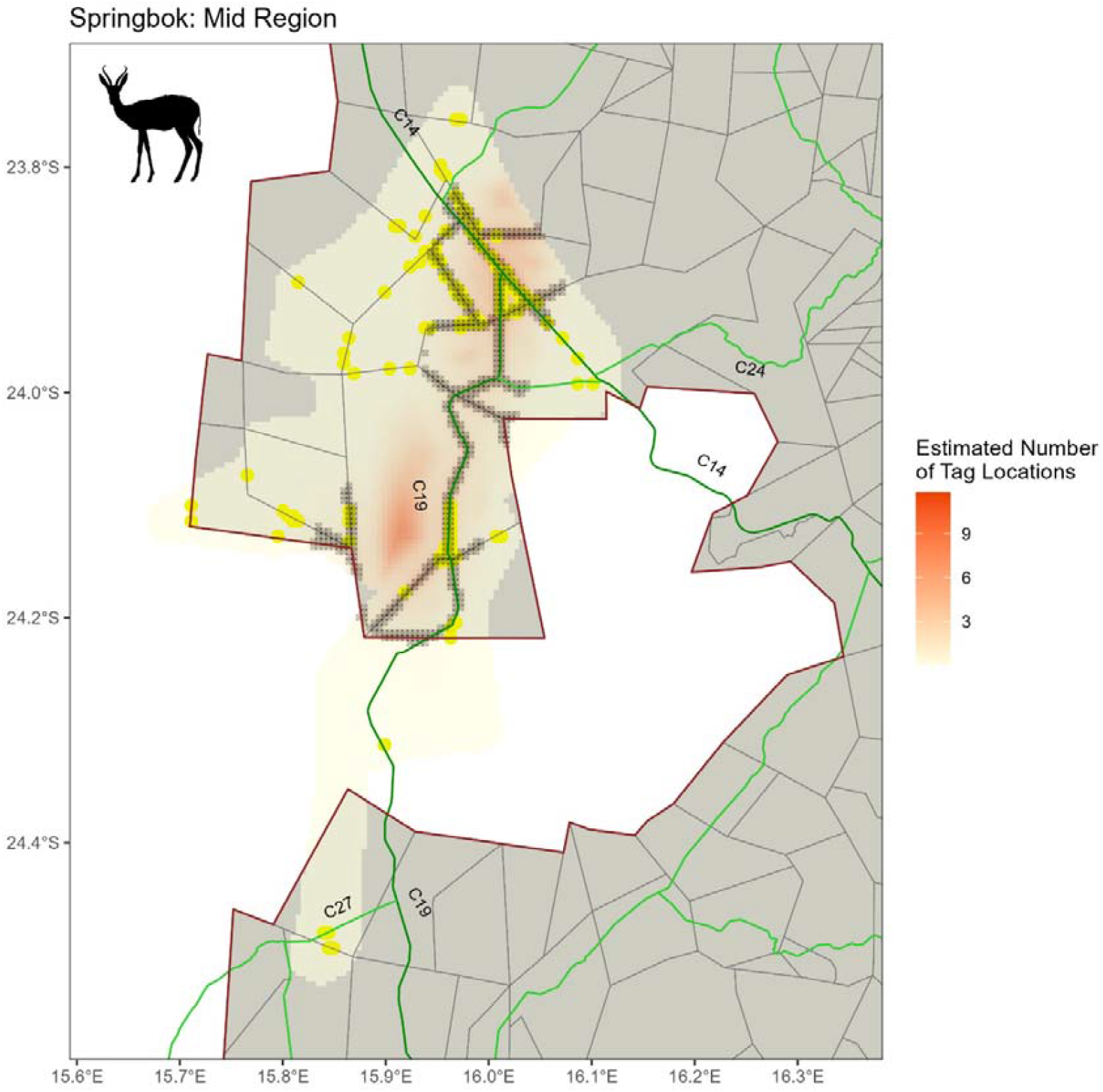
Estimated number of Springbok telemetry locations per cell. The ‘+’ symbols on the plots indicate where there are significantly more telemetry locations within 1 km of the barrier than would be expected under the universal barrier relationship. Grey lines: farm boundary, red line: NNP boundary, dark green lines: main roads, lime green lines: district roads.

**Figure 10.**
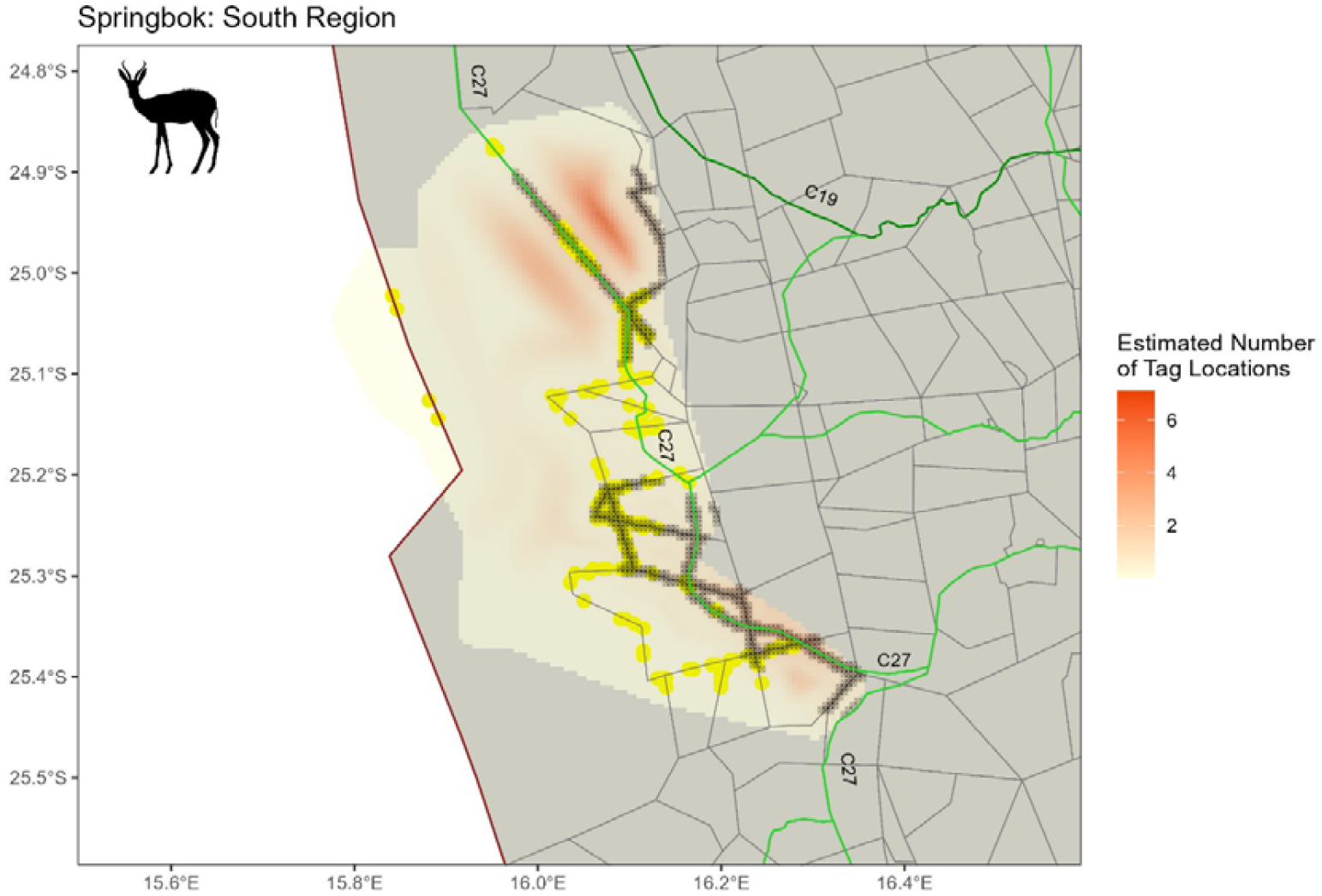
Estimated number of Springbok telemetry locations per cell. The ‘+’ symbols on the plots indicate where there are significantly more telemetry locations within 1 km of the barrier than would be expected under the universal barrier relationship. lines: farm boundary, red line: NNP boundary, dark green lines: main roads, lime green lines: district roads.

## Discussion

### Homerange sizes

Fences were found to affect homerange sizes and movement of all three species in the study area. The effect is however varied in intensity between species and per geographic location. Springbok and gemsbok were found to be most affected with several areas where fences significantly affected their movement, while Hartmann’s zebra were least affected and moved rather freely through most fences. This confirms the opinion of many farmers that the species break fences regularly.

Hartmann’s zebra in the study area ranged over exceptionally large areas, compared to those in the Daan Viljoen Game Reserve and the Etosha National Park (Joubert, 1973) where rainfall is somewhat higher than in the study area. They also exceeded the range sizes stated by Bothma & Du Toit (2016) for unrestricted areas, which is around 200 km^2^. The home ranges sizes (based on the 95% KDE) of six Hartmann’s mountain zebra monitored in the arid Kunene region of Namibia averaged 681 km^2^ and 256 km^2^ in the rainy and dry season respectively (Muntifering et al., 2019). The largest was approximately 950 km^2^, which is comparable to that of SAT1092 (925.50 km^2^) in our study area.

Hartmann’s zebra are said to be migratory (Gosling et al., 2019) and have separate dry and wet season ranges, some as far apart as 120 km (Joubert, 1973). Individuals in the Kunene region displayed differing degrees of overlap between dry and wet season ranges, with some having less pronounced (i.e. highly overlapping) seasonal home ranges (Muntifering et al., 2019). Animals in our study did not display a uniform migratory pattern, with individuals in the Naukluft and Nubib mountains generally having smaller and more concentrated home ranges, indicating that they were more sedentary, while those in the Central Namib and on farmland to the east of the GSNL generally displayed a distinctive north-south and east-west movement pattern respectively.

It is conceivable that the individuals with smaller and more concentrated home ranges selected areas close to permanent water sources, as was the case with Hartmann’s zebra in the Kunene region which did not venture more than 4 km from water sources (Muntifering *et al*. 2019). Contrary to findings by Joubert (1973) and Gosling et al. (2019), the collared individuals in this study did not follow seasonal patterns of movement, as each individual maintained a stable home range in different parts of the Naukluft mountains (north, east and west) over the entire 1.5 to 2 years that the three individuals were tracked.

Gemsbok in the Central Kalahari Game Reserve (CKGR), which were tracked by satellite over 9 months, had home ranges between 152.67 km^2^ to 1,154.63 km^2^, with a mean home range of 604.71±332.00 km^2^. This is similar to studies in the Central Kalahari Game Reserve (Skinner & Chimimba, 2005; Williamson & Williamson, 1984). They are however considerably smaller than home ranges in the Kgalagadi Transfrontier Park In the Kalahari Gemsbok National Park, radio-collared female MCP home ranges over the entire study period were between 430 km^2^ and 1,941 km^2^ (xL = 1,103 ± 629 km^2^, n=6), while they were between 2,529 km^2^ and 10,224 km^2^ (6,516 ± 3,867 km^2^, n=3) in the Gemsbok National Park (Knight, 2008). The larger home ranges of gemsbok individuals ranging in the south around the unfenced NamibRand Nature Reserve suggests a greater freedom of movement than those in the northern GSNL which had smaller home ranges. This was made especially clear by SAT1770’s extremely large home range (Figure 2), as well as that of SAT1107_1769, both of which ranged on the unfenced NamibRand and neighbouring Namib Naukluft Park.

Springbok home range and core area sizes were much larger compared to those reported in most literature. Most studies report about the territory sizes of springbok males (David, 1978; Jackson et al., 1993; Mason, 1976). Less is known about the home range sizes of females and of non-territorial males (Mechkour et al., 2008), especially in large and open areas such as the NamibRand Nature Reserve. The home ranges of springbok family herds (consisting of adult ewes, subadult female and male and female juveniles) are between 3.00 – 8.00 km^2^ in size (Furstenburg, 2006) while those of territorial males ranged anywhere between 0.01 – 0.70 km^2^ depending on the location in question (Apps & Smithers, 2012). Specifically, territory sizes were between 0.27 to 0.70 km^2^ in the former 31 km^2^ Jack Scott Nature Reserve in the Transvaal, South Africa (Mason, 1976), and 0.10 – 0.40 km^2^ in the 27.86 km^2^ Bontebok National Park, Swellendam, South Africa (David, 1978).

### Movements in relation to fences

The western part of the study area was less fragmented (mid region at about 15 km from barriers for the former and 10 km for the latter two) than the farmland in the east of the mid and south regions (mid region approximately 5 km for both regions). The main ecological consequences of roads and fences in the GSNL were related to barrier avoidance or blockage to movement, which influenced the movement patterns of ungulate species, and the physical effect of barriers, which prevented movement of individuals across these barriers. The presence of barriers also affected ungulate home range sizes, as those individuals ranging in more open systems (less barriers) had larger home ranges and vice versa.

In general, ungulate home range sizes were smaller in areas where there were physical barriers, including either roads (fenced and unfenced) or fences around land units. The effect of roads and fences on decreasing home range sizes has been found in tortoises (Peaden et al., 2017) and ungulates (Gulsby et al., 2011). The decrease in home range size could be due to multiple factors, including road avoidance behaviour, the inability to cross roads due to the physical barrier of fenced corridors, or the increased availability of food resources along roads that reduces the need of animals to move large distances to find forage (Gonser et al., 2009; Keken et al., 2019; Lightfoot & Whitford, 1991).

Fences and roadside fences are problematic for ungulate species, particularly in arid ecosystems, where they need to be highly mobile to access important food and water resources, and even more so in the face of climate change (Bennett et al., n.d.; Durant et al., 2017; Mbaiwa & Mbaiwa, 2006). In the GSNL, gemsbok and springbok movements were found to be more characteristic of movements performed by nomadic desert-dwelling ungulates that inhabit arid areas (Jonzén et al., 2011). As these are not spatially/temporally predictable and a lack of seasonality exists (Nandintsetseg et al., 2019), the highly unpredictable food availability coupled with the low food abundance in these environments does necessitate a high mobility of desert-dwelling ungulates in order to satisfy their food requirements. This high mobility of ungulates in the GSNL was reflected in their generally large home ranges. In those areas such as NamibRand, where the ungulate species were free to roam without major obstructions caused by fences, home ranges were generally larger and location points were less concentrated. Those individuals that ranged in areas with higher fence and road densities had considerably smaller home ranges, which is likely to have had a restrictive impact on their ability to move to areas with greater food abundance and quality after localised rainfall.

Fences and roads are more likely to negatively impact Hartmann’s zebra’s ability to move in response to their food and water requirements, as they display migratory movement patterns (Gosling et al., 2019). However, some individuals in this study that roamed on farmlands east to the GSNL had elongated home ranges, that were east-west. This indicates that their movement is not as affected by farm fences as much as gemsbok and springbok are. Alternatively, fences may not be as well maintained in mountainous areas due to their inaccessibility, and hence movement across these permeable barriers was possible. It is notable that Hartmann’s zebra home ranges were still much larger in areas without fences (such as the Central Namib area) than those with fences, indicating a degree of movement limitation caused by fences.

The availability of more forage along roads may have had an impact on decreasing home ranges of animals, especially for Hartmann’s zebra and springbok which were found to prefer areas close to barriers (within 1 km).

### Barriers which affected movements

One of the main causes of road-related movement limitations for ungulates in the GSNL are associated with physical hindrances (as most roads are delineated by fences on both sides). In general, the main roads C14 and C19 that run from the coast to Maltahöhe and from Solitaire to Maltahöhe respectively were the most significant barriers to the movements of all three species. Fences adjacent to district roads, including the C24 that runs from Rehoboth and meets the C14 around Remhoogte, the C27 that runs from NamibRand further south to Helmeringhausen, and the D0872 to Sossusvlei also constituted significant barriers. The D0855 leading past Neuras towards the C14 also significantly affected zebra movements. However, the identification of movement restrictions was based on only 40 sampled animals in a very large area – more problematic barriers might be identified with additional animal tracking data.

This study showed that the national park fence in the north region of the study area (Central Namib) was relatively impenetrable to Hartmann’s zebra movements, especially the portion between the C28 to the north and the D1982 to the south. Conversely, it did not totally obstruct ungulate movements in the mid (Naukluft area) and south (along NamibRand) where permeability permitted fence crossings. Farm (livestock) fences affected wildlife movements to varying degrees. While Hartmann’s zebra were able to cross some fences, springbok and gemsbok were not as successful, their movements sometimes being completely restricted within farms or along fences until they found a fence gap to cross. Hartmann’s zebra in the Kunene region similarly moved alongside the veterinary fence to find locations at which to cross (Muntifering et al., 2019).

Another important finding is that even when individuals succeeded in crossing fence segments, it often took a great amount of time for them to find weak spots, indicated by their persistent movement along farm and road segments. This movement barrier was particularly evident in a gemsbok (SAT1107_1769), which was tracked for the longest time (1,416 days). It moved along approximately 38 km of road fence and 21 km of farm fence in order to move eastwards (approximately 16 months of its tracking period) yet it never succeeded in crossing these barriers.

## Conclusion and Recommendations

To survive in the hyper-arid environment of the ProNamib, springbok, gemsbok and Hartmann’s mountain zebra need to move in order to access sproadic resources and surface water. In the GSNL, there is appetite to lobby landowners to remove fences in order to reduce fence mortalities, allow for unrestricted ungulate movement and improve the aesthetics for tourism.

Our study found that all three species, but particularly gemsbok and springbok are often restricted by fencing in the landscape. There are however specific fences which were found to be more restrictive to ungulate movements. Most of them are fences along roads, such as the C14 and C19 roads that run from the coast to Maltahöhe and from Solitaire to Maltahöhe. The fenceline along the C24 that runs from Rehoboth and meets the C14 around Remhoogte, the C27 that runs from NamibRand further south to Helmeringhausen, and the D0872 to Sossusvlei also constituted significant barriers to springbok and gemsbok specifically. The fenceline along the D0855 leading past Neuras towards the C14 significantly affected Hartmann’s zebra movements.

Our study cannot make a blanket recommendation to immediately remove all the fences which are shown to restrict the ungulate movements, since some of them are intended to reduce the risk of road collisions between vehicles and wildlife/livestock. Despite this, removal of some fences with low traffic volumes will be possible if combined with a reduced speed limit and warning signs. This has already been done on the C27 through NamibRand and does not seem to have resulted in an increase in vehicle-wildlife collisions. Where fencing is required to prevent off-road driving in conservation areas, removal of fence strands, or fixed fence-gap sections could be considered.

## Acknowledgements

We extend our gratitude to the Greater Sossusvlei Namib Landscape for initiating and supporting this study, as well as the Ministry of Environment Forestry and Tourism for conducting the capture and collaring operation as part of their mandate to reduce anthropogenic impacts on wildlife. We further thank Murray Tindall and Lee Tindall from the GSNL for logistical support and guidance, the Namibian Chamber of Environment for financial assistance and the Oppenheimer Generations Fellowship in People and Wildlife.

