## Supplementary Material for "What is the cost of that fence? The impact of fences on the movements of ungulates in a hyper-arid landscape"

**Results**

**Home rages**

| Species | Region | Sex | ID | Tag Locations | Home Range (km^2^) | Home Range 95% CI (km^2^) | Core Area (km^2^) | Core Area 95% CI (km^2^) |
| --- | --- | --- | --- | --- | --- | --- | --- | --- |
| Oryx | Mid | Female | SAT1101 | 1756 | 68.75 | (63.99, 76.26) | 24.75 | (21.99, 28.75) |
| Oryx | Mid | Female | SAT1102 | 1549 | 192 | (168.99, 220.76) | 74.75 | (56.99, 94.01) |
| Oryx | Mid | Female | SAT1103 | 1565 | 111.75 | (106.75, 122.01) | 42 | (38.75, 46.25) |
| Oryx | Mid | Female | SAT1104 | 1709 | 150.5 | (139.49, 166.26) | 49 | (44.24, 54.76) |
| Oryx | Mid | Female | SAT1105 | 1525 | 162.25 | (154.75, 181.51) | 64.25 | (60, 70.5) |
| Oryx | Mid | Female | SAT1106 | 1551 | 33.5 | (32.25, 35.5) | 12.75 | (12, 13.75) |
| Oryx | Mid | Female | SAT1765 | 2628 | 129.75 | (122.49, 138.25) | 44 | (40.75, 48) |
| Oryx | Mid | Female | SAT1766 | 2133 | 281 | (264, 307.26) | 82.75 | (75.75, 91.76) |
| Oryx | South | Female | SAT1107_1769 | 5064 | 704 | (142.47, 2437.28) | 204.75 | (30.73, 1312.42) |
| Oryx | South | Female | SAT1108 | 1581 | 224.75 | (47.98, 561.76) | 77 | (9.74, 305.29) |
| Oryx | South | Female | SAT1109 | 1506 | 476.75 | (442.75, 522) | 156.5 | (146.75, 171.51) |
| Oryx | South | Female | SAT1768 | 2473 | 434.75 | (397.99, 505.26) | 132.25 | (123.99, 147.76) |
| Oryx | South | Female | SAT1770 | 2486 | 1294 | (1062.49, 1483) | 365.75 | (255.74, 560.56) |
| Springbok | Mid | Male | SAT131 | 687 | 331 | (33.25, 841.51) | 102.25 | (8.75, 466.77) |
| Springbok | Mid | Male | st2010-2800 | 64 | 4 | (0, 76.5) | 2 | (0, 42.5) |
| Springbok | Mid | Male | st2010-2801 | 2528 | 167.5 | (31, 513.04) | 45 | (8, 277.25) |
| Springbok | Mid | Male | st2010-2803 | 2514 | 46.75 | (5.5, 218.52) | 9.25 | (1.5, 121.25) |
| Springbok | South | Female | st2010-2799 | 710 | 163.5 | (148.25, 182.01) | 53.25 | (49.5, 59.51) |
| Springbok | South | Male | SAT132.2 | 360 | 86.5 | (79.49, 96.5) | 25.5 | (22.5, 30.75) |
| Springbok | South | Male | SAT132 | 219 | 49.5 | (41.25, 66.51) | 16.25 | (12.75, 23.75) |
| Springbok | South | Male | SAT133 | 402 | 15.5 | (14.5, 18.26) | 5.25 | (5, 6.25) |
| Springbok | South | Male | SAT134 | 1161 | 258.5 | (13.74, 1158.5) | 71 | (7.49, 643.5) |
| Springbok | South | Male | SAT135 | 629 | 76.25 | (71, 91.01) | 26.5 | (24, 32) |
| Springbok | South | Male | st2010-2797 | 1375 | 220.5 | (201.75, 260) | 71 | (65.5, 82.26) |
| Springbok | South | Male | st2010-2798 | 593 | 289.25 | (230.74, 377.01) | 102.25 | (66, 156.53) |
| Zebra | Mid | Female | SAT1099 | 1424 | 44.5 | (23.75, 77.01) | 17.25 | (6.24, 40.01) |
| Zebra | Mid | Male | SAT1094 | 1728 | 64 | (58.25, 75.25) | 20.25 | (18.75, 23) |
| Zebra | Mid | Male | SAT1095 | 1518 | 128.75 | (117, 177.02) | 51.25 | (43.5, 65) |
| Zebra | Mid | Male | SAT1771 | 8898 | 154.5 | (53.22, 301.25) | 57.25 | (12.74, 148.26) |
| Zebra | Mid | Male | SAT1772 | 1845 | 185.25 | (34.99, 518.86) | 32 | (9.25, 285.57) |
| Zebra | Mid | Male | SAT1775 | 1743 | 135.25 | (121, 184.25) | 54.25 | (41.75, 73) |
| Zebra | North | Female | SAT1092 | 2150 | 1051.75 | (988.96, 1113.52) | 368.25 | (327.74, 411.51) |
| Zebra | North | Female | SAT1093 | 1743 | 484.75 | (461.74, 523.76) | 159.75 | (154, 170) |
| Zebra | North | Male | SAT1097 | 1753 | 565.75 | (513.25, 661.06) | 182.5 | (170.5, 206.26) |
| Zebra | North | Male | SAT1098 | 1746 | 365.75 | (340.99, 413.75) | 101 | (94.25, 113.51) |
| Zebra | South | Female | SAT1100 | 1451 | 61.75 | (58, 69) | 20.75 | (19.5, 23) |
| Zebra | South | Female | SAT1774 | 1675 | 139 | (118.99, 170.25) | 42.25 | (36.24, 51.25) |
| Zebra | South | Male | SAT1096 | 1531 | 80.75 | (75.25, 97.26) | 28.75 | (27.5, 31.25) |
| Zebra | South | Male | SAT1773 | 390 | 249 | (229.49, 271.26) | 81.25 | (73.75, 91.75) |
| Zebra | South | Male | SAT1776 | 1736 | 119.5 | (112.5, 129.25) | 32.25 | (29.5, 36.25) |

**Wildlife movements in relation to barriers**

*Hartmann’s mountain zebra*

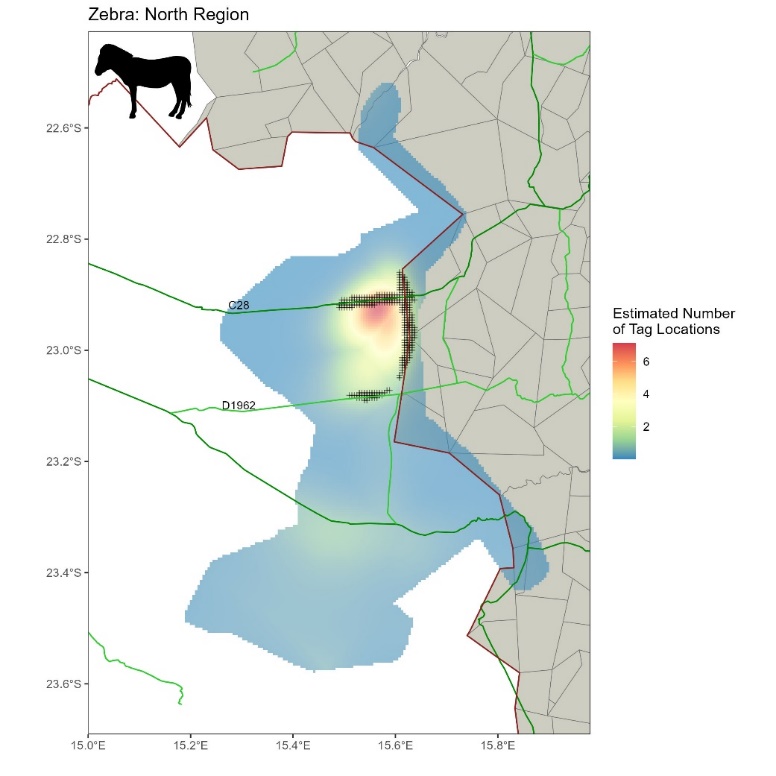

*Figure XX: Estimated number of Hartmann’s zebra telemetry locations per cell. The ‘+’ symbols on the plots indicate where there are significantly more telemetry locations within 1 km of the barrier than would be expected under the universal barrier relationship. Grey lines: farm boundary, red line: NNP boundary, dark green lines: main roads, lime green lines: district roads.*

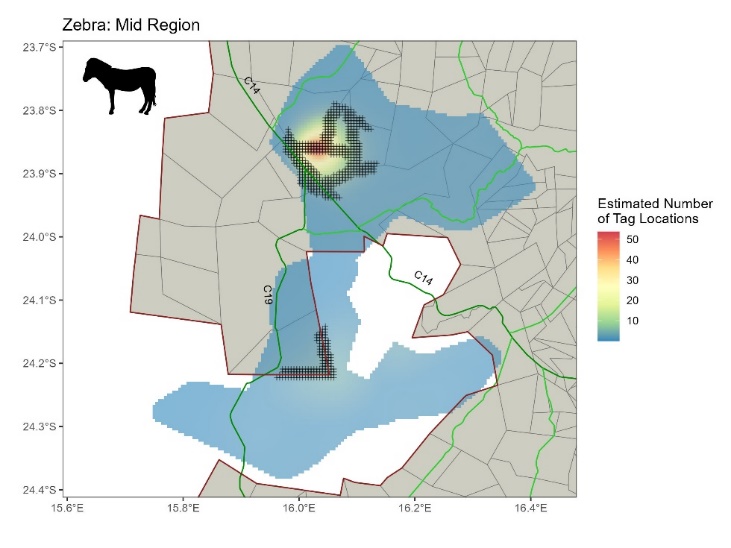

*Figure XX: Estimated number of Hartmann’s zebra telemetry locations per cell. The ‘+’ symbols on the plots indicate where there are significantly more telemetry locations within 1 km of the barrier than would be expected under the universal barrier relationship. Grey lines: farm boundary, red line: NNP boundary, dark green lines: main roads, lime green lines: district roads.*

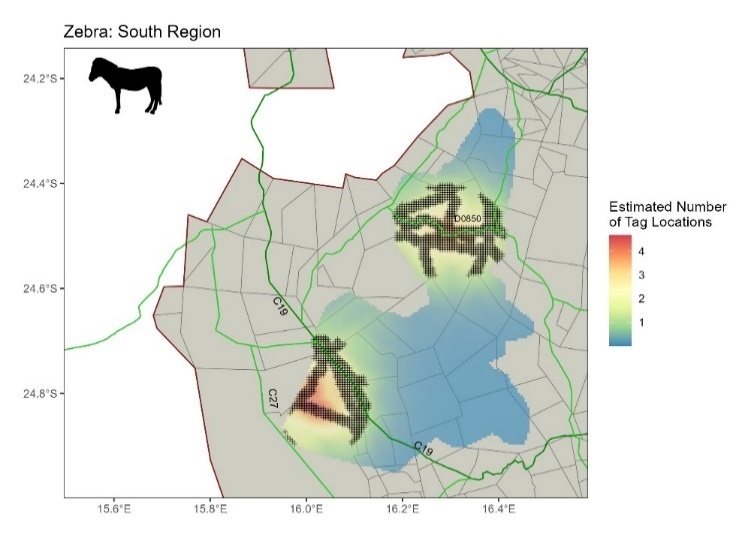

*Figure XX: Estimated number of Hartmann’s zebra telemetry locations per cell. The ‘+’ symbols on the plots indicate where there are significantly more telemetry locations within 1 km of the barrier than would be expected under the universal barrier relationship. Grey lines: farm boundary, red line: NNP boundary, dark green lines: main roads, lime green lines: district roads.*

*Gemsbok*

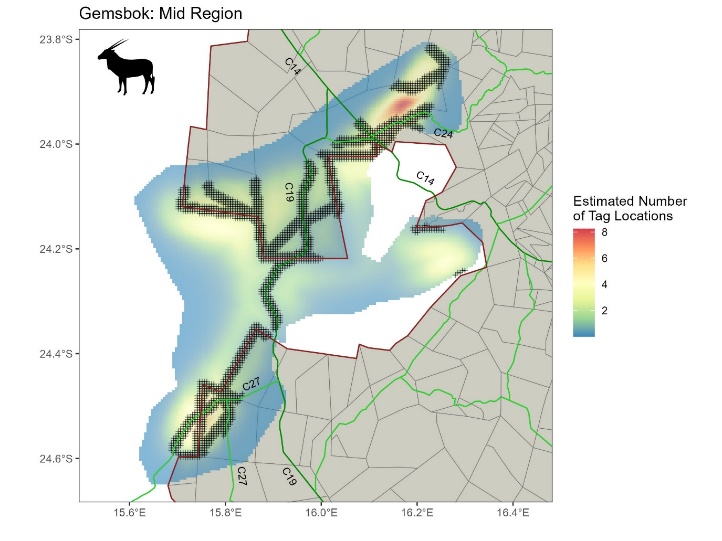

*Figure XX: Estimated number of Gemsbok telemetry locations per cell. The ‘+’ symbols on the plots indicate where there are significantly more telemetry locations within 1 km of the barrier than would be expected under the universal barrier relationship. Grey lines: farm boundary, red line: NNP boundary, dark green lines: main roads, lime green lines: district roads.*

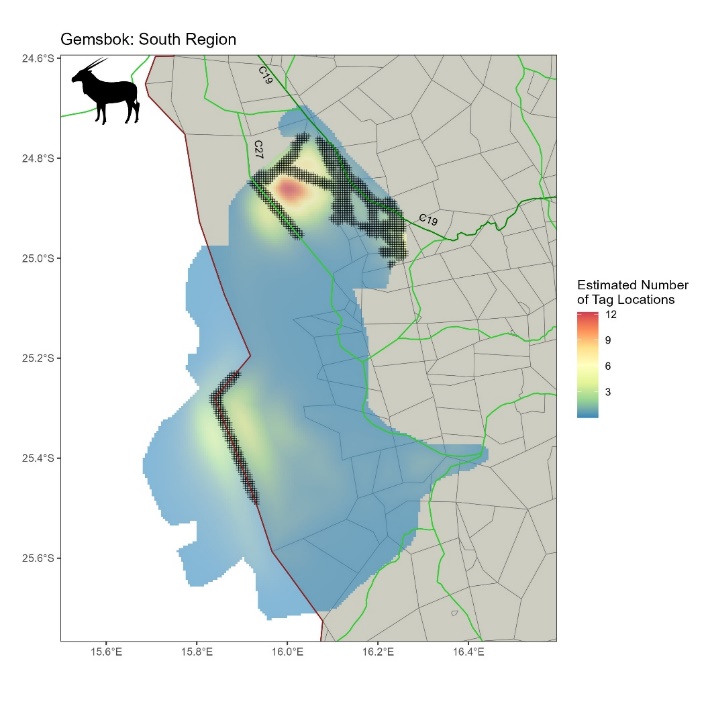

*Figure XX: Estimated number of Gemsbok telemetry locations per cell. The ‘+’ symbols on the plots indicate where there are significantly more telemetry locations within 1 km of the barrier than would be expected under the universal barrier relationship. Grey lines: farm boundary, red line: NNP boundary, dark green lines: main roads, lime green lines: district roads.*

*Springbok*

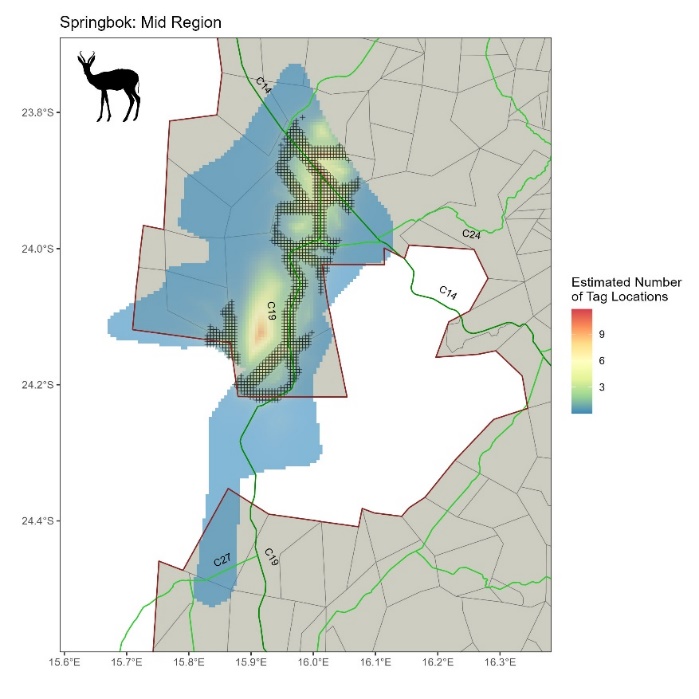

*Figure XX: Estimated number of Springbok telemetry locations per cell. The ‘+’ symbols on the plots indicate where there are significantly more telemetry locations within 1 km of the barrier than would be expected under the universal barrier relationship. Grey lines: farm boundary, red line: NNP boundary, dark green lines: main roads, lime green lines: district roads.*

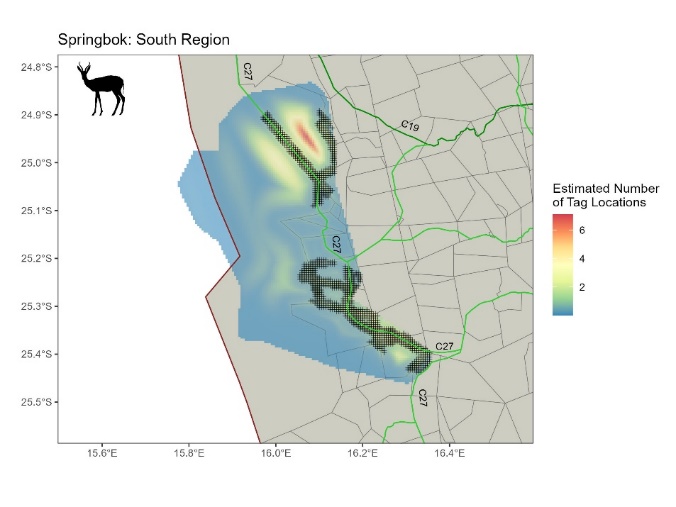

*Figure XX: Estimated number of Springbok telemetry locations per cell. The ‘+’ symbols on the plots indicate where there are significantly more telemetry locations within 1 km of the barrier than would be expected under the universal barrier relationship. Grey lines: farm boundary, red line: NNP boundary, dark green lines: main roads, lime green lines: district roads.*
